# The geometry of low- and high-level perceptual spaces

**DOI:** 10.1101/2023.09.02.556032

**Authors:** Suniyya A. Waraich, Jonathan D. Victor

## Abstract

Low-level features are typically continuous (e.g., the gamut between two colors), but semantic information is often categorical (there is no corresponding gradient between dog and turtle) and hierarchical (animals live in land, water, or air). To determine the impact of these differences on cognitive representations, we characterized the geometry of perceptual spaces of five domains: a domain dominated by semantic information (animal names presented as words), a domain dominated by low-level features (colored textures), and three intermediate domains (animal images, lightly-texturized animal images that were easy to recognize, and heavily-texturized animal images that were difficult to recognize). Each domain had 37 stimuli derived from the same animal names. From 13 subjects (9F), we gathered similarity judgments in each domain via an efficient psychophysical ranking paradigm. We then built geometric models of each domain for each subject, in which distances between stimuli accounted for subjects’ similarity judgments and intrinsic uncertainty. Remarkably, the five domains had similar global properties: each required 5 to 7 dimensions, and a modest amount of spherical curvature provided the best fit. However, the arrangement of the stimuli within these embeddings depended on the level of semantic information: dendrograms derived from semantic domains (word, image, and lightly texturized images) were more ‘tree-like’ than those from feature-dominated domains (heavily texturized images and textures). Thus, the perceptual spaces of domains along this feature-dominated to semantic-dominated gradient have a similar global geometry, but the points within the spaces shift to a tree-like organization when semantic information dominates.

**Significance Statement:** Understanding the nature of knowledge representation is a fundamental goal of systems neuroscience. Low-level visual features (e.g., color), form continuous domains, while semantic information is typically organized into categories and subcategories. Here, using a novel psychophysical paradigm and computational modeling strategy, we find that despite these major differences, the mental representations of these domains lie in spaces with similar overall geometry. However, within these spaces, semantic information is arranged in a more tree-like representation, and the transition to tree-like representations is relatively abrupt once semantic information becomes apparent. These findings provide insight into visual stream processing at an algorithmic level. Furthermore, they support the idea that processing along the ventral stream reflects commonalities of intrinsic cortical function.

## Introduction

Low-level features and objects constitute qualitatively different kinds of domains, and this suggests that their cognitive representations may have different characteristics. Whereas low-level features typically form continuous domains, objects appear to form categories and hierarchies (Kemp and Tenenbaum, 2008). For example, between any two colors, such as blue and green, there is a continuous gamut that can be achieved by mixing. This gamut has no natural parallel for a pair of generic objects, e.g., a pencil and a fox, and objects naturally form hierarchical categories (inanimate vs. animate; land vs. sea vs. air). One might therefore expect that the mental representations of these two kinds of information are fundamentally different, and that the geometry of their perceptual spaces (Kriegeskorte and Kievit, 2013) will mirror these differences.

Psychophysical studies provide partial support for this idea (Zaidi, 2013). Color and visual texture can both be described by low-dimensional perceptual spaces that are approximately Euclidean (Maxwell, 1860; Chubb et al., 2004; Victor et al., 2015, 2017); semantic spaces typically require many more dimensions (Huth et al., 2012; Hebart et al., 2020, 2023). Studies of representation of objects (Kriegeskorte et al., 2008; Connolly et al., 2012; King et al., 2019; Hebart et al., 2020) show that while continuous spatial models can describe the perceptual spaces of both low-level and semantic information, representation of semantic information can also be explained by hierarchical, tree-like models (Sattath and Tversky, 1977; Kiani et al., 2007; Kriegeskorte et al., 2008; Connolly et al., 2012; Dehaqani et al., 2016). However, it is unclear whether these differences in representational geometry are merely quantitative (e.g., the precise number of dimensions required or the extent to which the representational space is Euclidean), or more fundamental (continuous domains vs. hierarchical categories).

Neurophysiologic and computational studies also do not provide a clear answer. On one hand, brain regions that represent different kinds of information are distinct: low-level features are represented in early visual areas, but object information is primarily represented in inferotemporal cortex (Martin et al., 1996). Moreover, functionally specialized regions in the brain respond to specific kinds of higher-level visual stimuli (e.g., faces, places, bodies, and food) (Kanwisher et al., 1997; Jain et al., 2023). These areas (FFA, PPA, EBA etc.) vary in their cytoarchitectural details (Weiner et al., 2017), suggesting that their neural computations and representational formats are distinct. On the other hand, despite these functional differences, there are striking commonalities across neocortex: lamination (Van Essen and Maunsell, 1983), columnar organization (Mountcastle, 1997), and circuit motifs – e.g., the canonical microcircuit (Douglas and Martin, 1991; Bastos et al., 2012). These commonalities suggest that differences in how information is represented may be superficial, reflecting differences in inputs rather than representational formats. Deep neural network models of the ventral visual stream (Kriegeskorte, 2015; Cichy et al., 2016) – able to represent both continuous and hierarchical information (Saxe et al., 2019; St-Yves et al., 2023) – appear to support this view. Often, these models (Kriegeskorte, 2015; Yamins and DiCarlo, 2016; King et al., 2019) have at least some similarly structured layers (e.g., multiple convolutional layers, or multiple fully connected layers), but nevertheless show the shift from representing low-level features to representing objects, that parallels the human ventral stream.

Here, we directly compare the representations of low-level features and semantic information, via a parallel analysis of five stimulus domains with varying levels of low-level and semantic content. At one extreme, we study a domain of visual textures; at the other, we study domains of animal names and images of animals. The two intermediate domains are lightly-texturized and heavily-texturized images of animals. In each domain, we collect similarity judgments via the same psychophysical paradigm, construct the participants’ perceptual spaces, and compare their geometric characteristics. As we show, these characteristics change systematically as semantic content increases, but the change is best captured by geometric characteristics that are more subtle than dimension *per se*.

## Materials and Methods

### Participants

13 participants, recruited from personnel at Weill Cornell Medicine and the surrounding institutions, participated in the study (9F, 4M, age range 23 to 62 years). All participants had normal color vision and had 20/20 visual acuity after correction. Color vision was tested by an online Red-Green color-blindness test (https://www.colorlitelens.com/color-blindness-test.html). All participants were fluent in English. Most were native English speakers (S1, S4, S6, S8, S9, S10, S11, S12 and S13); the others grew up with English as their second language. They provided informed consent following institutional guidelines and the Declaration of Helsinki, and the protocol was approved by the Institutional Review Board of Weill Cornell Medical College. Three participants (S1, S5 and S7) were part of the laboratory and included one of the authors (S5: SAW), while the remaining ten were naïve to the purpose of the experiments. 10 participants participated in all experiments for all five domains; the exceptions were S6 (image-like and image domains), S9 (texture, texture-like and word domains) and S13 (texture-like, image, and word domains). Additionally, four participants (S1, S5, S6 and S8), participated in at least one of the two control-domain experiments.

### Procedure and Apparatus

The experimental paradigm, along with stimulus-creation and analysis scripts, are detailed in (Waraich and Victor, 2022) and are summarized here.

Participants responded to self-paced trials (described below) via mouse-clicks on stimuli displayed on a computer monitor. For in-laboratory experiments (11 participants), the display was a 13-inch laptop screen (MacBook Pro with Retina Display, resolution 2560×1600, Color Profile: LG-TV), viewed binocularly from a distance of 57 cm. The display had a mean luminance of 55 cd/m^2^. Data were collected via Zoom for two participants (S1 and S7) during the pandemic via screen-share and remote control of the mouse. These participants calibrated their distance from their respective screens themselves with a tape measure to maintain the same visual angles as the other participants. At this distance the visual angle of the display was 12.2 degrees, from the left edge of the leftmost stimulus to the right edge of the rightmost stimulus, and the visual angle of each stimulus was typically 2.25 degrees. In the texture and texture-like domains, the size of each check within the stimulus was approximately 13.3 arcmin. For the word experiment, the visual display was slightly larger, approximately 15 degrees, to ensure adequate spacing between longer words.

Stimuli were presented using experiments created with the open-source PsychoPy software, following editing of images in Adobe Illustrator. Experiment scripts and most analysis scripts were written in Python and run with Python version 3.7 or 3.8. Scripts for the clustering analysis were written and run in MATLAB.

### Software Accessibility

Code for running experiments and the Euclidean model-fitting is available on GitHub (https://github.com/jvlab/similarities) and described in (Waraich and Victor, 2022). Additional code will be provided upon acceptance and publication.

## Stimuli

### Overview

Motivated by the hypothesis that semantics are represented in a hierarchical and categorical manner that contrasts with a continuous representation of low-level features, we chose familiar animals that naturally lend themselves to categorization (Connolly et al., 2012) as the basis for the study. From the chosen set of animals, we created five stimulus domains as detailed below, with varying degrees of semantic information and low-level features (see Figure 1). At one extreme were domains consisting of animal names (the “word” domain), and animal pictures (the “image” domain). At the other extreme was the “texture” domain, in which small tiles derived from the image were scrambled. In between we created two intermediate domains, one more texture-like and the other image-like.

**1.**
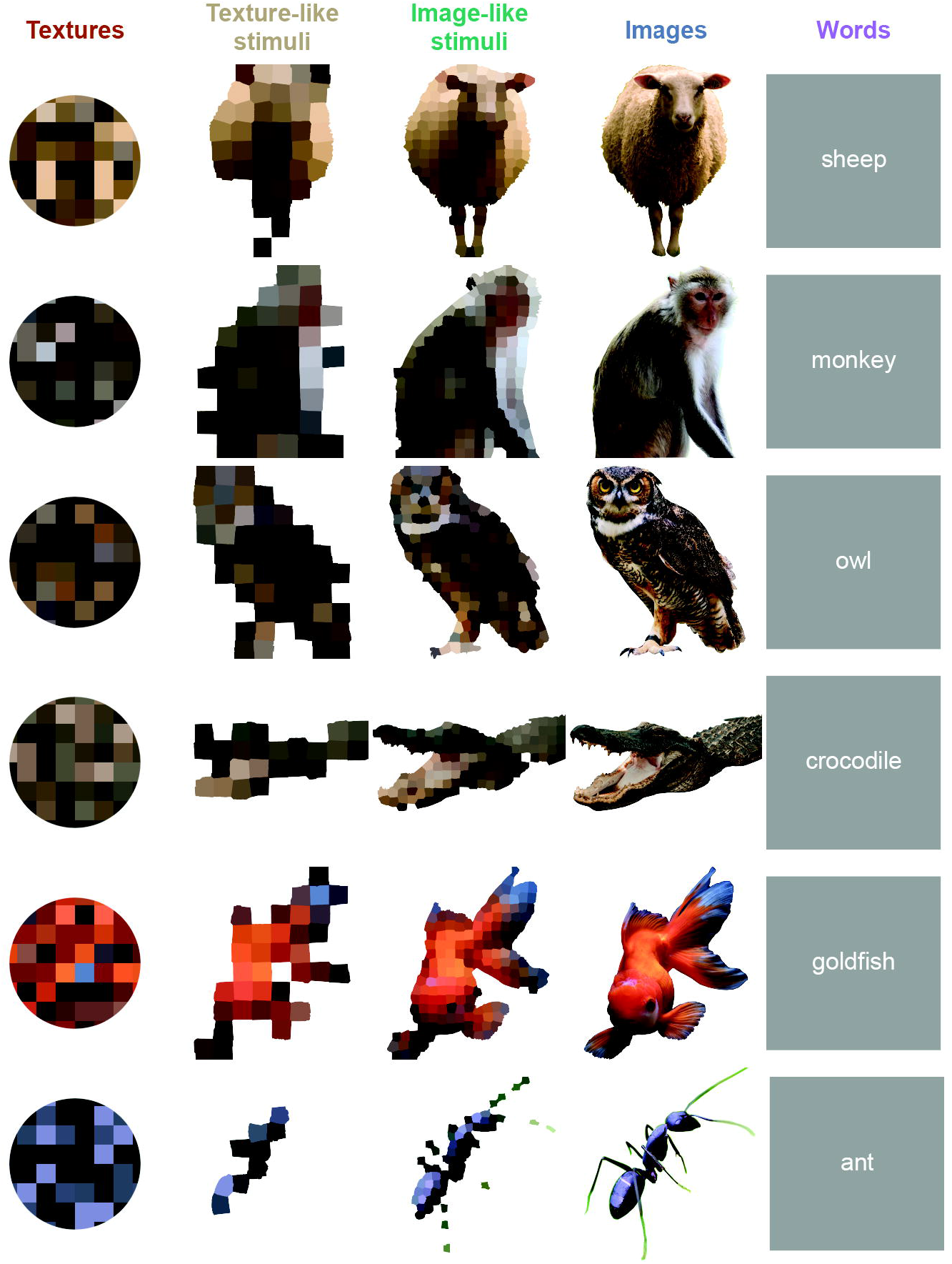
Example stimuli. The six rows show corresponding stimuli from the five domains (columns): textures, texture-like stimuli, image-like stimuli, images, and words. The full set of 37 animals used for each stimulus domain included mammals, reptiles, birds, and insects.

### Word Domain

A set of 37 animal names were chosen from the top 50 animal name terms returned by WordNet in 2019. WordNet’s Popularity Index was used as a proxy for familiarity. Selections were made to include a range of categories (e.g., land vs. sea vs. air), sizes, and colors. Word stimuli were presented in white text in lowercase text against a gray background, in Arial font, and subtended 1.1 to 3.1 deg depending on the length of the word, (the shortest being ‘owl’/ ‘bat’ and the longest being ‘crocodile’).

### Image Domain

The image domain was comprised of naturalistic, colorful pictures of the animals. For each animal, one representative, readily recognizable image was downloaded from https://unsplash.com/, an open library of high-resolution images. Most images were a front-facing view but for some birds and insects, we chose side or top-down views since these were more readily recognized.

Stimuli were created from these images by background removal (after manual tracing), followed by cropping and resizing to 4000 by 4000 pixels. A small number of images that were initially at slightly lower resolution (e.g., rat, bluebird) were up-sampled using bicubic interpolation via Python’s PIL package. Animals’ body sizes were approximately constant across the final set of images, but necessarily varied somewhat because of shape differences. The size of the presented images was approximately 2.25 by 2.25 deg, after down-sampling by the stimulus presentation software, PsychoPy.

### Texture Domain

Texture stimuli retained the approximate color and luminance distribution of the image of the corresponding animal, but no shape information. Each texture stimulus was created by choosing 100 RGB values randomly from the corresponding texture-like stimulus and then using them to color checks within a grid of 10 x 10 checks. A circular mask of diameter 1.6 deg was then applied to the grid. This number was chosen such that the total area of the texture patches was equal to the average area of the animal pictures.

### Texture-like and Image-like Domains

These domains retained partial shape information (e.g., a rough outline of the animals), but this information was degraded (more degraded for the texture-like domain, less for the image-like domain) using ‘super-pixels’. Super-pixels were made by applying Simple Linear Iterative Clustering (SLIC) (Achanta et al., 2012) to each of the corresponding images in the image domain. Briefly, SLIC clusters groups of pixels in an image into a given number of patches called superpixels based on 1) how close the pixels are to each other, and 2) how similar the colors of the pixels are. The distance between pixels is a weighted average of the above two metrics. The clustering determined by SLIC depends on three parameters (Achanta et al., 2012):i)threshold – which we kept constant at 0.002, ii) n_segments (number of segments), which determined the number of ‘super-pixels’ and iii) compactness, which controlled the relative weight to assign color or distance when assigning pixels to clusters. The compactness parameter indirectly controlled the shape of the super-pixels, making them more compact and squarish for higher values.

Our goal was to convert the original images into texture-forms composed of squarish checks of about the same size, so that the finer details were obscured, and the animals were hard to recognize (the texture-like domain) or easy to recognize (the image-like domain). The values of n_segments and compactness needed to do this depended on the area taken up by the figure (larger figures required lower values for n_segments and higher values for compactness), so we selected parameters for the five largest and five smallest stimuli by eye, and then interpolated these values as a linear function of the stimulus area to determine parameters for the remaining stimuli. This procedure was carried out separately for the texture-like domain to create images in which recognizability was difficult to impossible – but shape information was still present – and the image-like domain, in which recognizability was easier but not immediate. Image-like stimuli were similar to texture-like stimuli but with smaller super-pixels.

For the texture-like domain experiment, for the first two participants (S1 and S7), a single exemplar of each animal was used. Because these participants reported that they recognized some of the individual animals, for the subsequent participants, we created 15 exemplars per animal, and randomly selected an exemplar on every trial. We did the same in the image-like domain experiment, for all participants. Exemplars were created by jittering SLIC parameters. For the other domains, single exemplars were used throughout.

### Control Domains

Control-domain stimuli probed the three-parameter domain of color and the domain of spatially uncorrelated grayscale textures, which is also known to be three-dimensional (Chubb et al., 2004). These were derived from the texture-domain stimuli. For each animal, the corresponding color stimulus consisted of a disk (diameter 1.6 deg) with the mean RGB value of the corresponding texture-domain stimulus. Likewise, for each animal, a grayscale texture was created by converting its texture-domain stimulus to grayscale (with 256 gray levels) using Python’s PIL package.

## Experimental Design

### Paradigm

The experiments probed participants’ judgments of relative similarity within each domain. In each experiment, stimuli were chosen from a single domain and grouped into 222 unique trials, consisting of 8 “comparison” stimuli and a 9^th^ “reference” stimulus in the center of the display (see Figure 2A for an example trial from the word domain). Each of these trials was repeated 5 times, over the course of 10 sessions of approximately 1 hour each, conducted on separate days. In the 222 unique trials, each of the 37 stimuli appeared as the reference stimulus in 6 trials, and each reference stimulus was paired with each comparison stimulus at least once.

**2.**
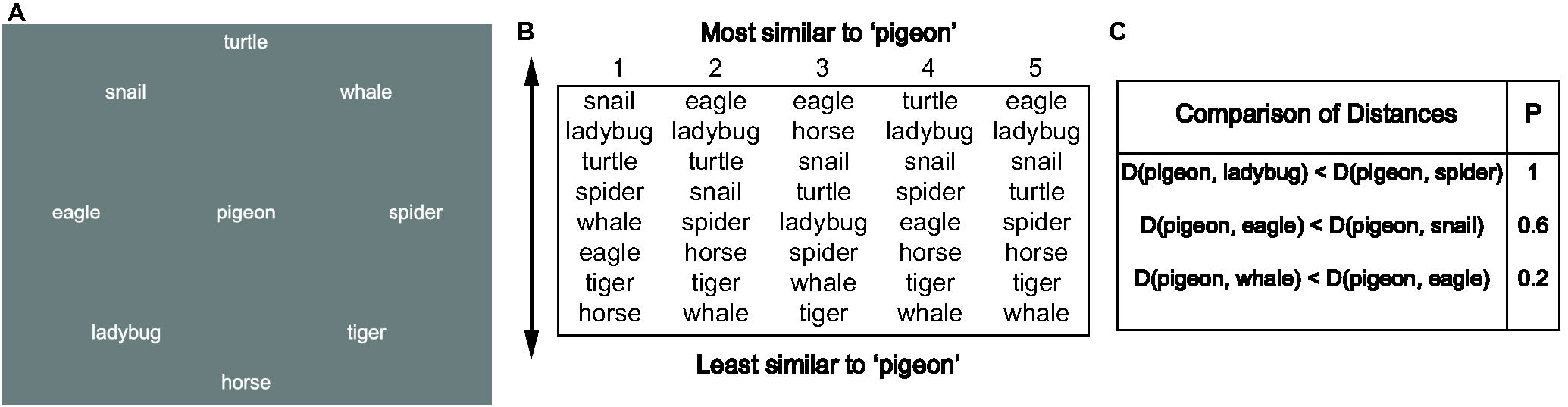
Collection of similarity judgments. **A:** a sample trial from the word experiment showing the reference (‘pigeon’) surrounded by eight comparison stimuli. **B:** judgments ranking similarity of the comparison stimuli relative to the reference are obtained in five separate presentations of this stimulus subset (interleaved among presentations of many other combinations). **C:** judgments are pooled and interpreted as choice probabilities.

### Task

At the start of the study, participants were informed that they would view a different domain of stimuli in each set of ten sessions, and that in all domains, the stimuli were derived from the same set of 37 animals. The task, which was explained orally, was to rank the eight comparison stimuli in order of similarity to the reference, by clicking the most similar stimulus to the reference first and least similar stimulus last. These instructions also appeared onscreen at the start of every session. Participants could not change their judgments within a trial; as participants clicked stimuli, the stimuli faded. No instructions were given regarding how to gauge the similarity between stimuli. Trials were self-paced, and participants were encouraged to take breaks if needed. See Figure 2B for an example of participant responses for the 5 repeats of an example trial. At the conclusion of participation, participants were debriefed and asked to describe their strategy for determining similarity. Most participants indicated that they used a combination of color and pattern in the texture domain; color, shape and size in the texture-like and image-like domains; and additionally, size and various category classifications in the image-like and image domains. In the word domain, participants mostly used size and various category classifications; for exceptions, see below (“Exclusion Criteria”).

### Randomization

The sequence of the five experiments was pseudo-randomly chosen to reduce the possibility that an easy-to-recognize object would prime a more difficult one: the image domain was only presented after the image-like domain, which was only presented after the texture-like domain. To avoid priming words with an image, the word domain was presented prior to the image experiment. No sequence constraints were placed on the texture experiment as guessing the identity of the animal from a texture was not very likely. These constraints resulted in 15 different possible orders; they were randomly assigned to participants.

The grouping of stimuli into trials was designed to assay the effect of context, i.e., the stimuli not explicitly involved in a comparison between the reference stimulus and a pair of comparison stimuli. We used two strategies for this grouping: a pseudo-randomized scheme (S1-S9), and a scheme intended to accentuate context effects (S10-S13). These are detailed in the section “Assessing Context Effects.” Since we found that the effect of context was minimal, these cohorts were analyzed together.

Other than for two participants in the texture domain (S1 and S5), all participants within a cohort were shown the same trials. The trial order was randomized across participants within each pair of sessions (which contained the full set of 222 unique trials). Within each trial, the position of the 8 surrounding stimuli was also randomized across repetitions and participants.

### Modeling the Perceptual Spaces

To characterize the participants’ latent mental representations of the stimulus domains, we used the framework of perceptual spaces (Zaidi, 2013): for each domain, we attempted to identify “coordinates” corresponding to each of the 37 stimuli, so that the perceived dissimilarities between the stimuli – as calculated from distances between the stimulus coordinates – would account for the similarity judgments.

Coordinates were determined using a variant of non-metric multidimensional scaling developed for this study, detailed in (Waraich and Victor, 2022). This algorithm recognizes that participants’ judgments provided rankings of distances (rather than numerical distance estimates), but also makes use of the fact that these judgments are repeated, yielding choice probabilities. The algorithm then seeks an assignment of coordinates to each stimulus, for which the Euclidean distances between stimulus points best account for a participant’s set of judgments.

### Preprocessing

The first step in the analysis is to convert the rank judgments into choice probabilities for each triad, i.e., the fraction of times that a participant judged a reference stimulus to be more similar to one comparison stimulus than another (see Fig 2C). Since each trial has 8 comparison stimuli, a single trial provides information on 28 (=8×7/2) such triads. We assume participants rank the stimuli by making these pairwise comparisons independently, choosing the stimulus with the smallest distance (perceived dissimilarity) to the reference at each click. Thus, an experiment (222 unique trials) yields 6216 (=28 × 222) pairwise comparisons, each of which are repeated 5 or 10 times over the full experiment. Typical triads appear in just one unique trial, so its choice probabilities can have the values 0, 0.2, 0.4, 0.6, 0.8 and 1. 222 of the triads appear in two unique trials (to assay context effects); their choice probabilities can have the values 0, 0.1, …, 0.9, and 1.0.

### Model-Fitting Procedure and Benchmarks

#### Overview of the Model-Fitting Procedure

The modeling procedure (Waraich and Victor, 2022) seeks a set of coordinates between stimulus “points” that maximizes the log-likelihood of the observed choice probabilities. Separate runs of the algorithm are carried out for each participant, each stimulus domain, and, for model spaces of 1, 2, 3, 4, 5, 6, and 7 dimensions.

The algorithm is initialized by a heuristic that assigns large distances to stimulus pairs that are most often judged to be most dis-similar, and then uses gradient descent to adjust the coordinates until the cost function (the negative log-likelihood of the observed set of responses) is minimized. The algorithm terminates when the cost function reaches a minimum or after an iteration limit (typically 50,000) is reached. Full details on the fitting procedure are reported in (Waraich and Victor, 2022).

#### Decision Model

The decision model assumes that uncertainty arises when distances (i.e., dissimilarities) are compared, and that there is no uncertainty in the coordinates, or in the computation of the distances from their coordinates. Specifically, we assume that the perceptual distance between two stimuli corresponds to the standard Euclidean distance between their coordinates, and that, when two distances are compared, the difference in distances is corrupted by the addition of a Gaussian noise of standard deviation (*σ*). Thus, the probability of choosing one distance, *d(s*_*r*,_*s*_*m*_) as greater than another, *d(s*_*r*,_*s*_*n*_) is modeled by the following as in (Maloney and Joong, 2003; Victor et al., 2017):

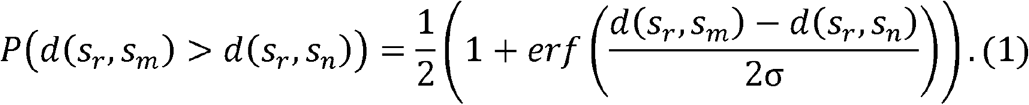

#### Cost Function

The cost function is the negative log-likelihood of the observed set of judgments, given the above model for distances and distance comparisons. Each triad of stimuli that appear in a trial together, *r*,*i*,*j* with *i* < *j* contributes a separate term:

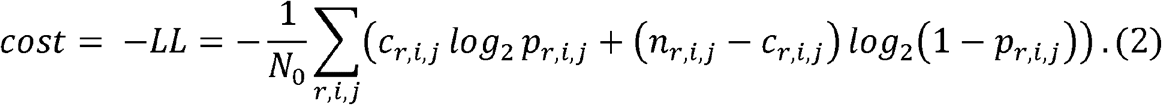

*N*_0_ is the total number of triad judgments (222 trials x 5 repetitions x 28 triads per trial=31,080 total triad judgments). *n*_*r*,*i*,*j*_ is the number of times each triad appears over all sessions, and is either 5 or 10. *p*_*r*,*i*,*j*_, is the modeled choice probability, as determined from eq. (1):

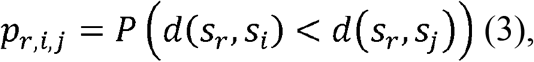

*c*_*r*,*i*,*j*_ counts the number of times *i* is chosen to be more similar to *r* than *j* is to *r*:

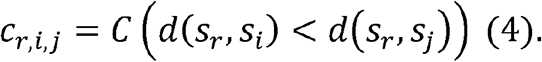

As is typical of multidimensional scaling procedures, the model fit is unchanged if all of the coordinates are rotated or translated, as this does not change their distances. For this reason, our analyses only consider the intrinsic geometry of the coordinates that minimize the cost.

However, in contrast to standard multidimensional scaling procedures in which the goodness of fit is unchanged if all distances are multiplied by a constant factor, here, the absolute sizes of the distances can be interpreted. The reason is that here, the choice probabilities depend on the ratio of distances to the noise parameter *σ* (eq. 1). We therefore fix *σ*=1, and as a result, the absolute distances and coordinates are in units of “decision noise.”

#### Comparing Model Performance Against Benchmarks

To help interpret the log-likelihood of a coordinate model, we compare its log-likelihood to a lower and an upper bound. The lower bound model (the ‘random’ model) posits that all choice probabilities are equal to 0.5. The upper bound model (the ‘geometrically unconstrained’ model) assumes that for each comparison the model probability, *p*_*r*,*i*,*j*_ is exactly equal to the empirical choice probability, *c*_*r*,*i*,*j*_*/n*_*r*,*i*,*j*_. It ignores constraints stemming from relationships between choice probabilities and distances, and merely recognizes that, even if a model accounts for the choice probabilities exactly, it cannot predict each decision. The ‘geometrically unconstrained’ model has an overfitting bias because it exactly matches the observed choice probabilities.

We estimated this overfitting bias via simulations, in which the ground-truth probabilities were known. Points corresponding to the stimuli were sampled from a simulated ‘perceptual space’ (for details, see ‘Validations’), and ground-truth choice probabilities were determined from eq. 1. Then, we simulated experimental datasets by Bernoulli sampling of these ground-truth choice probabilities. Next, we used the choice probabilities from the simulated datasets to compute the log-likelihood of the geometrically unconstrained model (LL in eq. 2). Additionally, we computed the log-likelihood of the ground-truth model (in which the choice probabilities are determined from the simulated coordinates), which would be unknown in a real experiment. The difference between these log-likelihoods – between the geometrically unconstrained model and the ground-truth model – is then an estimate of the overfitting bias.

Before implementing this procedure, we first assessed what factors affected the size of this bias. We sampled points from three kinds of ‘perceptual spaces’: 1) a *D*-dimensional Gaussian distribution, 2) uniformly from the surface of a sphere in a *D*-dimensional Euclidean space, or 3) uniformly from the interior of a *D*-dimensional hypercube (where *D=*2,3,4,5). We used a range of values for the noise parameter, *σ*. Across these geometries, the bias depended on model dimension (because mean distances grew larger with model dimension) and the ratio of the root-mean-square distance to *σ*, but was independent of the sampling strategy. We therefore ran further simulations with points sampled from a *D*-dimensional Gaussian distribution (repeated 80 times for dimensions 1 to 5 and 60 times for dimensions 6 and 7) to establish empirical bias correction, based on the ratio of the root-mean-squared (RMS) distance to *σ* . To apply this bias correction to a model, we increased the raw model log-likelihood by the estimated median bias (given by the ratio of RMS distance to *σ* of that model).

### Extension to Non-Euclidean Models

We extended the above procedure to model perceptual spaces with two kinds of non-Euclidean geometries: hyperbolic (shown (Zhou et al., 2018) to be a good model for certain olfactory spaces) and spherical. These geometries have contrasting characteristics: hyperbolic space has a uniform negative curvature (like an infinite saddle); spherical space has a uniform positive curvature (like the surface of a sphere). A related contrast between these geometries is that in a hyperbolic space, the volume that is within a given distance *r* of a point grows exponentially (i.e.,*V*(*r*) ∼*e*^*kr*^, while in spherical space, volume has a finite limit: the entire sphere. In a Euclidean space, in which curvature is zero, volume grows as a power-law function of distance, i.e., *V*(*r*) ∼*r*^*D*^, where *D* is the dimension.

To determine how well these non-Euclidean spaces could account for our data, we introduced a curvature parameter into the above fitting procedure. As detailed below, the curvature parameter influenced how distance was calculated from the coordinates, but not how the resulting distances led to a choice probability (eq. 1). Zero curvature, the Euclidean case, along with the coordinates determined above, served as the starting point. We gradually moved the curvature parameter away from zero to either progressively more negative values (the hyperbolic case) or more positive values (the spherical case), adjusting the coordinate values at each step to minimize the cost (eq. 2). We refer to the curvature parameter as *λ* in the hyperbolic case and as *μ* in the spherical case.

In the hyperbolic case, the distance between two points *x*=(*x*_1_,…,*x*_*D*_) and *y*=(*y*_1_,…,*y*_*D*_) is calculated in two steps: first, by mapping each of the coordinate *D*-tuples into a *D*+1-tuple on a hyperboloid (the ‘Loid’ model), and second, by computing the distance between them in the ‘Loid’ model (Tabaghi and Dokmanić, 2020) (see Figure 3A). A point *z*=(*z*_0_,*z*_1_,…, *z*_*D*_) lies on the hyperboloid if it satisfies the equation:

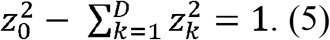

Following (Tabaghi and Dokmanić, 2020), the mapping *z=L*(*x*) from a *D*-tuple *x*=(*x*_1_,…,*x*_*D*_)to *D*+1a-tuple *z*=(z_0_,*z*_1_,…,*z*_*D*_) on the hyperboloid is

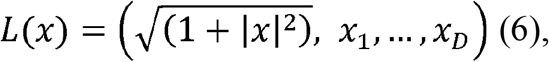

where 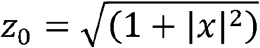and 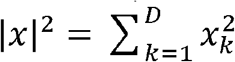.

**3.**
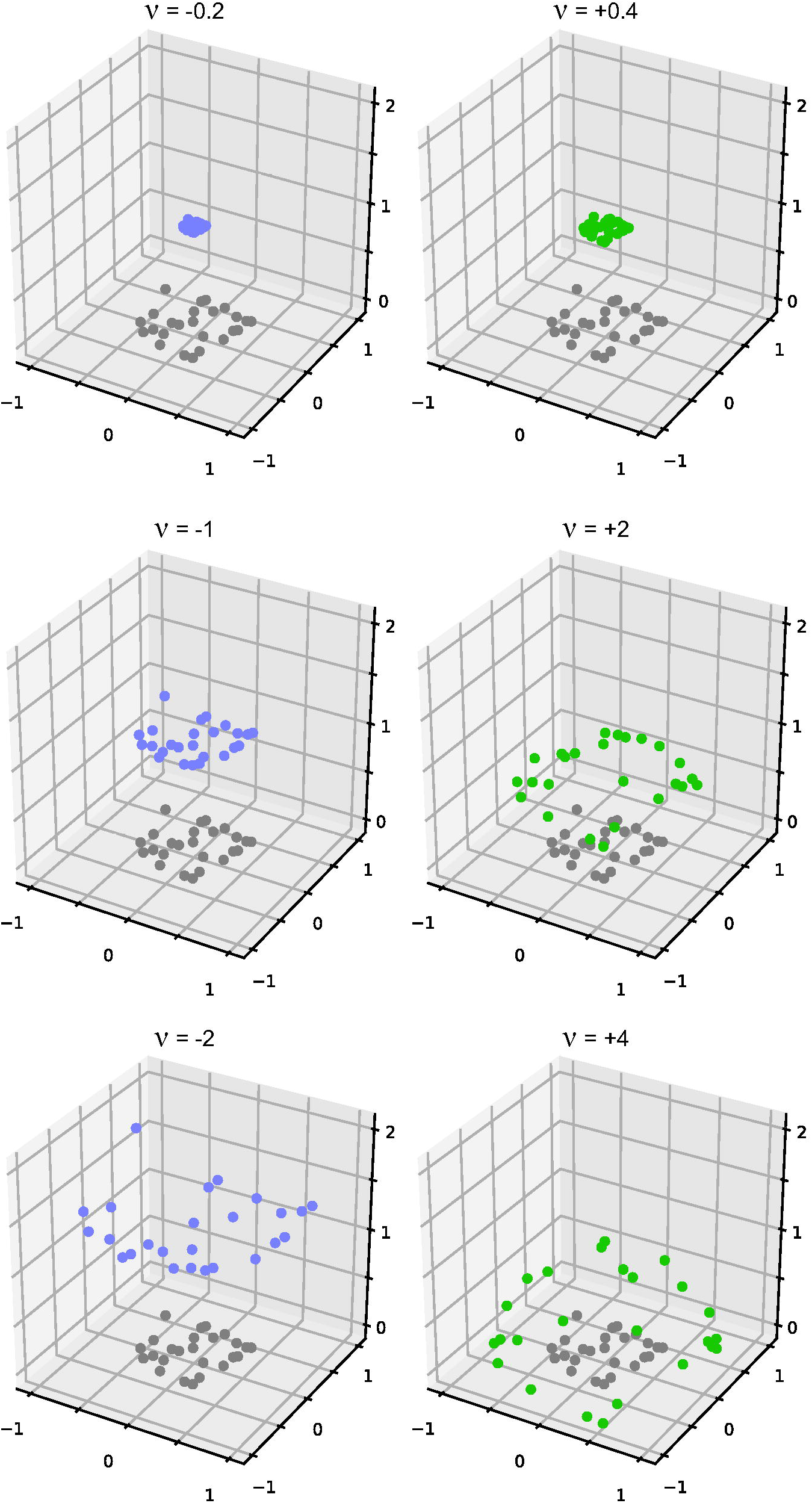
Strategy for assessing hyperbolic (left) or spherical (right) geometries. An initial model fit in a Euclidean space (here, in 2 dimensions, gray points) is projected onto a hyperboloid (purple) with a range of values of negative curvature (left) or onto a sphere with a range of values of positive curvature (right). Positions of the points are then adjusted so that the distances in the curved space best account for the observed choice probabilities. As detailed in the text, curvature is gradually increased, and at each step, the solution identified for the smaller curvature is used as the starting point for the larger curvature. For hyperbolic geometry, *v* = − *λ* (see text equation 7); for spherical geometry, *v* = 2*μ*(see text equation 10).

The distance on the hyperboloid is calculated in the standard fashion (Tabaghi and Dokmanić, 2020)

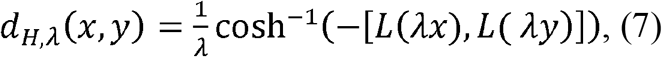

where the inner product between two points on the hyperboloid, *z* and *z*^*′*^ is defined by:

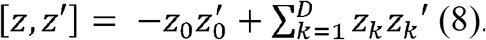

Note that when *λ*→0,*d*_*H*, *λ*_ approaches the ordinary Euclidean distance. Increasing values of *λ* correspond to increasingly negative curvatures, since the mapping *L*(*λx*) extends further and further into the hyperboloid.

The spherical case follows a parallel strategy but uses a stereographic, rather than an orthographic, projection from the *D*-tuple to the sphere |*z*|^2^in a *D +*1-dimensional space (Figure 3B). This mapping, *z*=*T*(*x*), is given by

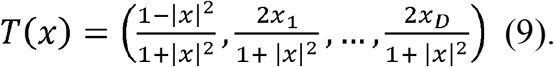

We use this stereographic mapping rather than an orthographic mapping (e.g., eq. (6) with the first coordinate replaced by 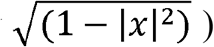 so that the domain of the mapping is unrestricted and its range covers both hemispheres (except for the point *z*=(-1,0,0,…,0) of the range, which is approached arbitrarily closely as *x* grows without bound).

On the sphere, distance is proportional to arc length, which is determined by the angle at the origin subtended by the points *T*(*μx*) and *T*(*μy*). Thus we use

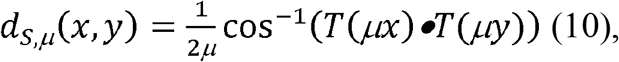

where • is the ordinary dot-product and *μ* controls the curvature. The initial factor 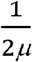 is chosen so that, parallel to the hyperbolic case, as *μ*→0,*d*_*S,μ*_ approaches the ordinary Euclidean distance.

To determine the curvature corresponding to this mapping, we determine the radius of the sphere corresponding to (10). To do this, we note that one-quarter of the length of a great circle is given by *d*_*S,μ*_(*x*,*y*) when *T*(*μx*) and *T*(*μy*) are orthogonal. From eq. (10), this distance is 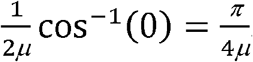. So a great circle has length 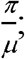 the sphere’s radius is 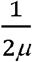, and the curvature is 2*μ*.

To put negative and positive curvature on a common footing, we use a single signed quantity,*v*, for both: *v=*−*λ* in the hyperbolic case, and *v=*2*μ* in the spherical case.

Since each of these models used, as their starting point, a model fit with a closely related geometry (the Euclidean geometry when curvature is near zero, or a similar curvature as the curvature moves away from zero), the maximum number of iterations used for adjusting the coordinates was 30,000 rather than 50,000. Increasing this parameter for datasets from a few participants did not make a substantial difference (<0.001) in the model log-likelihood.

### Validations

As an analytical control, we tested whether our modeling approach correctly recovers the dimension of simulated perceptual spaces. For the simulated spaces, we sampled 37 points from spaces of *D*_*true*_=2, 3, 4, 5, 6 and 7 dimensions and used their Euclidean distances to simulate psychophysical ranking judgments, using the decision model of eq. 1. We then applied the above multidimensional scaling procedure to these judgements for models of various dimensions *D*_*model*_, and compared the log likelihood of the model fits to the log likelihood of the ground-truth model (i.e., where the model probabilities are derived using eq. 1 from the true coordinate distances). For *D*_*model*_ < *D*_*true*_, but not when *D*_*model*_≥ *D*_*true*_, there was a gap between the ground-truth log-likelihood and the fitted model log-likelihood. These simulations were carried out three ways for each dimension, *D*_*true*_=2,3,4,5,6 and 7: sampling from Gaussian distributions, from the surface of a sphere and uniformly within a *D*-dimensional sphere, and with 10 simulations per condition.

As a control both for the analysis pipeline and the decision model, we tested whether the experimental protocol and analysis pipeline would correctly recover the dimensionality of visual perceptual spaces for which the dimension has been established by previous studies. To this end, we conducted two control psychophysical experiments, in which stimuli sampled perceptual spaces that are known to be 3-dimensional (Maxwell, 1860; Chubb et al., 2004): color space (S1, S5, S6) and the space of uncorrelated grayscale textures (S1, S5, S8). As expected, 3-dimensional models accounted for similarity judgments from the control domains.

As a test of whether an incorrect decision model might result in a spuriously high dimensionality, we carried out simulations in which an alternative decision model was used to generate the judgments, and then analyzed the data with our standard (incorrect) decision model. Specifically, when simulating judgments, this alternative decision model assumed there was noise at the level of coordinates (i.e., the coordinates were jittered from their true positions on each trial before any distances were calculated and compared). However, during fitting we kept the assumption that the noise was at the comparison stage instead. We ran simulations (16 times with 3 values of the noise parameter each) in which we sampled points from a 3-dimensional space. We found that the true dimensionality model could still be recovered, despite the incorrect decision model.

### Clustering Analyses

To further characterize the set of distances inferred by the analysis pipeline, we performed hierarchical clustering and then characterized the structure of the resulting tree.

For each participant and each domain, the distances inferred from the coordinate model were used to drive hierarchical clustering of the 37 stimuli, using the ‘linkage’ and ‘dendrogram’ functions in MATLAB. The calculations presented here reflect hierarchical clustering performed on the 5-dimensional Euclidean coordinate model; calculations based on other model dimensions and models with spherical curvature (see below) yielded the same pattern.

Once a tree was obtained for each participant and domain, we calculated two statistics to quantify the extent to which the tree resembled a perfectly balanced binary tree:

1. <u>Balance:</u> At each branch along the tree, we calculated the ratio of the size of the bigger subtree to the size of the smaller one. This ratio was then averaged across all branches. In a perfectly balanced, binary tree, we expect this average to be 1; for an unbalanced tree, it is larger
2. <u>Informativeness:</u> We calculate the cumulative information available at each level of the tree, as follows: H = −Σ*p*_*i*_ log_2_(*p*_*i*_), where *p*_*i*_ is the number of elements in each subtree, *i*, at a given level, divided by the total number of elements (37). This quantity is maximized by a binary tree, which yields *k* bits of information at level *k*. We present results for *k*=3, which is approximately halfway through the tree – since at the root and last level of the tree, information is necessarily constrained to be 0 and log_2_ 37 respectively

For comparisons across domains, we only included the participants who had completed all 5 experiments and excluded those participants (S2, S3 and S8) who had used idiosyncratic strategies (see Exclusion Criteria).

We report results utilizing MATLAB’s ‘single’ (nearest neighbor) linkage method. For other linkage methods (‘complete’, ‘centroid’, ‘average’, ‘median’ and Ward’), similar trends were observed. The largest difference between trees for the texture and word domains was found with the ‘single’ method.

### Exclusion Criteria

For comparison across domains, we omit the three participants who did not provide data on all five domains (S6, S9 and S13).

In some analyses, we distinguish 3 participants (S2, S3, S8), as they reported use of idiosyncratic strategies in the word experiment. Participants S2 and S8 both used one-dimensional metrics to rank the animals – intelligence (S2), and suitability as a pet (S8). S3’s word data is excluded from the group analysis because S3 reported basing these judgments on phonetic similarity or letters in common, ignoring word meaning.

### Context Effects

As mentioned above, each judgment of similarity of two comparison stimuli to a reference was made in the context of six other comparison stimuli that were not explicitly relevant to this judgment. We assayed the possible influence of this ‘context’ in several ways, each making use of the fact that in each of the experiments, 222 of the unique triads were presented in two different contexts. We refer to the trials in which these triads appeared as the ‘context’ trials.

For participants S1-S9, the stimuli that constituted the two contexts for the ‘context’ triads were chosen at random. We determined whether the difference in choice probabilities observed between the two contexts was greater than would be expected by chance. As detailed in (Waraich and Victor, 2022), this determination was made by computing an imbalance statistic. The imbalance statistic is a sum of contributions from each context triad, where each context triad contributes the negative log of the Fisher exact test probability for the 2×2 table of choices in the two contexts. We then determine the probability of observing this value under the null hypothesis that the choice probability of a given comparison was the same across contexts. Across the 9 participants and 5 domains (40 comparisons), we found marginal evidence for context effects in two instances: for S1 and S7 in the image domain (p < 0.05 after FDR-correction).

In participants S10-S13, we actively manipulated some of the configurations of the ‘context’ trials, in an attempt to maximize the context effect according to Tversky’s Diagnosticity Principle (Tversky, 1977; Evers and Lakens, 2014). Tversky’s Diagnosticity Principle states that if certain features (e.g., habitat or size in our case) have diagnostic value, they will play an enhanced role in similarity judgments because participants tend to group the stimuli into clusters based on shared features. To apply this idea, we generated ‘context’ trials in which, for a given triad, the reference belonged to one category (e.g., ‘land-dwelling’) and the two comparison stimuli belonged to two other categories (e.g., ‘water-dwelling/traveling’ and ‘air-traveling’ respectively), and generated contexts in which the context – the accompanying comparison stimuli -- were either entirely of one or the other of these categories (e.g., ‘water-dwelling/traveling’ and ‘air-traveling’). The Diagnosticity Principle predicts that participants will group the stimuli based on shared features, and that the sole stimulus left out of the larger cluster will be perceived as more like the reference than the stimulus in the larger cluster (Evers and Lakens, 2014). We implemented this with 13 pairs of ‘context’ trials based on the land/water/air categories, and 10 pairs of ‘context’ trials based on size (small/medium/large); in the latter case, the reference stimulus was always ‘medium.’

Across the four participants and the five domains, there was only one combination (S12, image-like) in which the context effect, as measured by the imbalance statistic, was significant (p <0.01 after FDR-correction).

The Diagnosticity Principle also predicts the direction of the context effect, and we tested whether this prediction held. For each context triad, we expected one comparison stimulus to be judged as closer to the reference in one context and further from the reference in the other. We then looked at the choice probabilities in all pairs of context trials in which there was a shift, and asked, by binomial statistics, were the directions of the shift in the directions predicted by Diagnosticity? Only one dataset confirmed the predictions of the Diagnosticity Principle (S11, word domain, p <0.05 after FDR-correction).

As none of the analyses revealed more than marginal evidence of context effects, all further analyses were carried on ‘context’ and non-’context’ trials combined.

## Results

### Overview

The aim of this study was to determine how the geometry of perceptual spaces depends on the extent to which they represent low-level features vs. semantic content. To address this, we asked participants to judge the similarity between stimuli within five domains, including domains with substantial semantic content (words and pictures of common animals), a domain with only low-level features (textures created by scrambling these images), and two intermediate domains. In each case, we determined the geometry of the perceptual space implied by the similarity judgments, and we compared the geometries in several ways.

As we detail below, we found that the dimensionality of these domains was comparable, but there was a systematic difference in the way that points (i.e., stimuli) were arranged: in the domains with a greater semantic weighting, the arrangement was more tree-like. The most substantial difference was observed between the texture-like and word domains, suggesting that perceptual spaces of stimuli become more hierarchical when they progress from early to late-stage domains.

### Dimensionality

To study the geometry of each perceptual domain, we analyzed the similarity judgments with a form of non-metric multidimensional scaling (see Methods). This procedure modeled the data by arranging the stimuli in a Euclidean space in a way in which the similarity judgments corresponded to distances between the points. Figure 4 shows the ability of these models to account for the judgments, in terms of how well they predict the participants’ responses. To aid in interpretation, the predictive power of these geometric models is compared to the maximum possible predictive accuracy (determined from a model that uses the participants’ observed choice probabilities, without any geometric constraints), and a baseline predictive accuracy (a model that posits that all choices are made randomly). As shown in Figure 4, in all cases, a sufficiently high-dimensional embedding of the stimuli provided a reasonably accurate account of the participants’ judgments.

**4.**
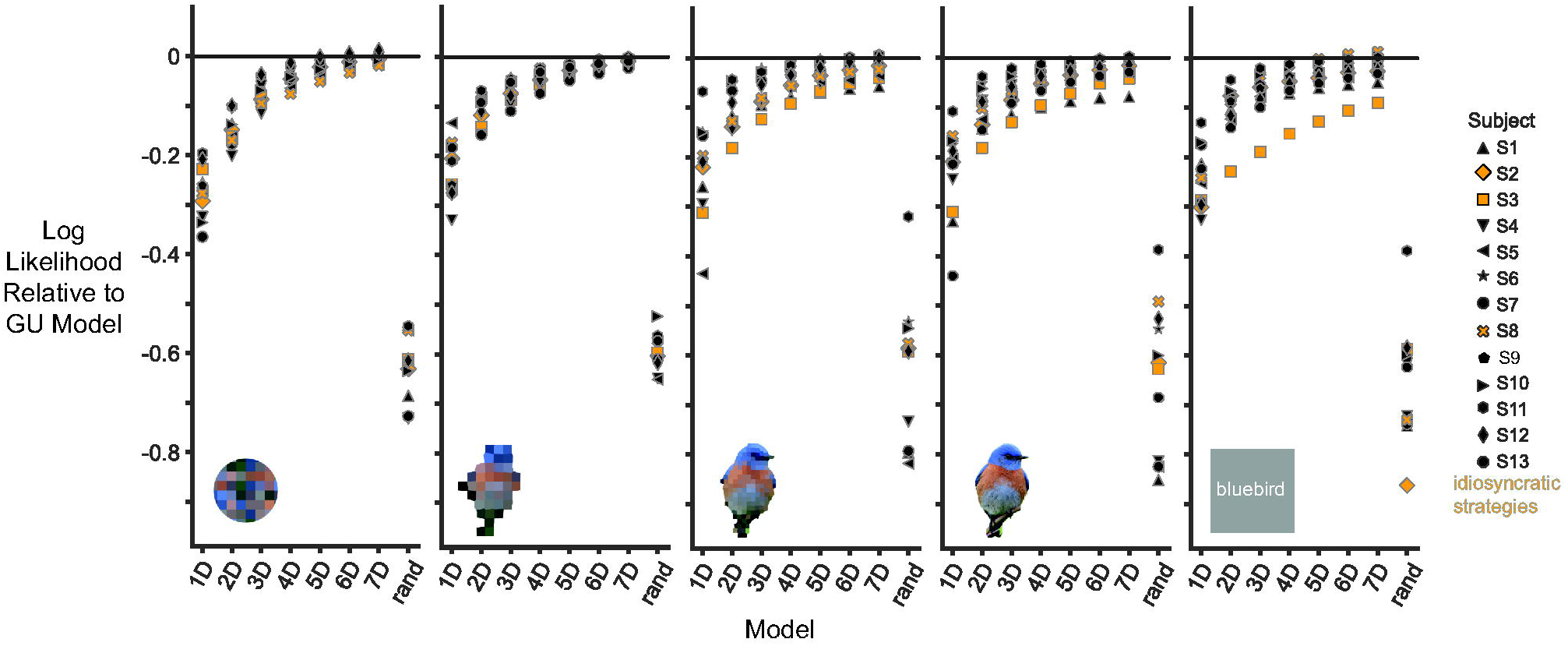
Dependence of model fit on number of dimensions for the five domains and all participants. The log-likelihood, per trial, is shown relative to the log likelihood of the geometrically unconstrained (GU) model, a model in which choice probabilities exactly match the data (with a correction for overfitting). ‘rand’ is a lower bound, the relative log likelihood of the observed responses, under the assumption that similarity judgments are random. Colored symbols indicate participants whose data were excluded from some group analyses (see ‘Exclusion Criteria’). The data from participants S4, S7 and S9, in the word domain were previously presented in (Waraich and Victor, 2022, Figure 5).

We estimated the number of dimensions needed to model each stimulus domain, by comparing how the log-likelihoods of models changed with the number of dimensions. As expected, increasing the number of dimensions from 1 to 7 improved the explanatory power of the model (see Figure 4). This trajectory showed a common pattern across domains, with a plateau in the log-likelihoods around dimensions 4 and 5. While there was some variability in the extent to which the plateau reached the level of a complete account, this variability was primarily participant-dependent, rather than domain-dependent. We thus concluded there was no clear difference in dimensionality across domains.

Additionally, considering the principal components of the obtained 7-dimensional models, we found that the first 5 principal components explained most of the variance (83.5-95.5%, mean=90.3%) in the coordinate positions, for all participants and domains. Thus, we use the coordinates from the 5-dimensional perceptual spaces in some analyses below that probe the geometry at a finer level of detail. The trends we report also hold for models taken from other dimensions.

While 5-dimensional models are almost as good as 6- or 7-dimensional models in their log-likelihoods, even 7-dimensional models fail to explain all datasets (Figure 4: the gap between the model log-likelihoods of the 7-dimensional models and the geometrically unconstrained model log-likelihood). An auxiliary analysis (see Methods: Validation) showed that this unexplained gap was not likely to be due to the use of an incorrect noise model.

### Do the perceptual spaces of early and late-stage domains vary in curvature?

The surprisingly high dimensionality needed to account for the data in a Euclidean model raises the possibility that a non-Euclidean perceptual space might account for the data with fewer dimensions. That is, we hypothesized that perhaps the perceptual spaces were curved, and a lower-dimensional curved space would better fit the similarity judgments better than a Euclidean (flat) space. Positive curvature corresponds to spherical spaces, e.g., in two dimensions, the surface of a 3-dimensional sphere, whereas negative curvature corresponds to hyperbolic space. An important consequence of curvature is the volume of neighborhoods around a point in the resulting space. For positively curved, spherical spaces, the volume reaches a maximum as distance from the origin, *r*, increases; in a *D*-dimensional Euclidean space, this volume scales with *r*^*D*^; in a hyperbolic space, it increases more rapidly, growing exponentially with respect to *r*. Furthermore, a hyperbolic geometry is an important alternative to consider because it has been linked to olfactory perceptual spaces (Zhou et al., 2018) and because hyperbolic spaces are a natural way to accommodate trees, which have been used to describe mental representations of objects (Kriegeskorte et al., 2008; Connolly et al., 2012).

To test the hypothesis that a non-Euclidean space might be a better model of the perceptual spaces, we extended the geometric modeling procedure to allow for negative or positive curvature. We then determined whether a curved space would provide a better fit than a Euclidean space by parametrically varying the curvature away from zero, keeping the number of dimensions fixed. As shown in Figure 5 (data shown for 2-dimensional fits (A), similar results for 7-dimensional fits (B)), the quality of the fit declined for large values of either negative or positive curvature – with a more rapid decline for the latter, likely because in spaces with positive curvature, the maximum distance is bounded. Across all domains and all participants, the best fit was for an approximately Euclidean space with a modest amount of spherical (positive) curvature, typically *v* = 0.2 . We will interpret this value below.

**5.**
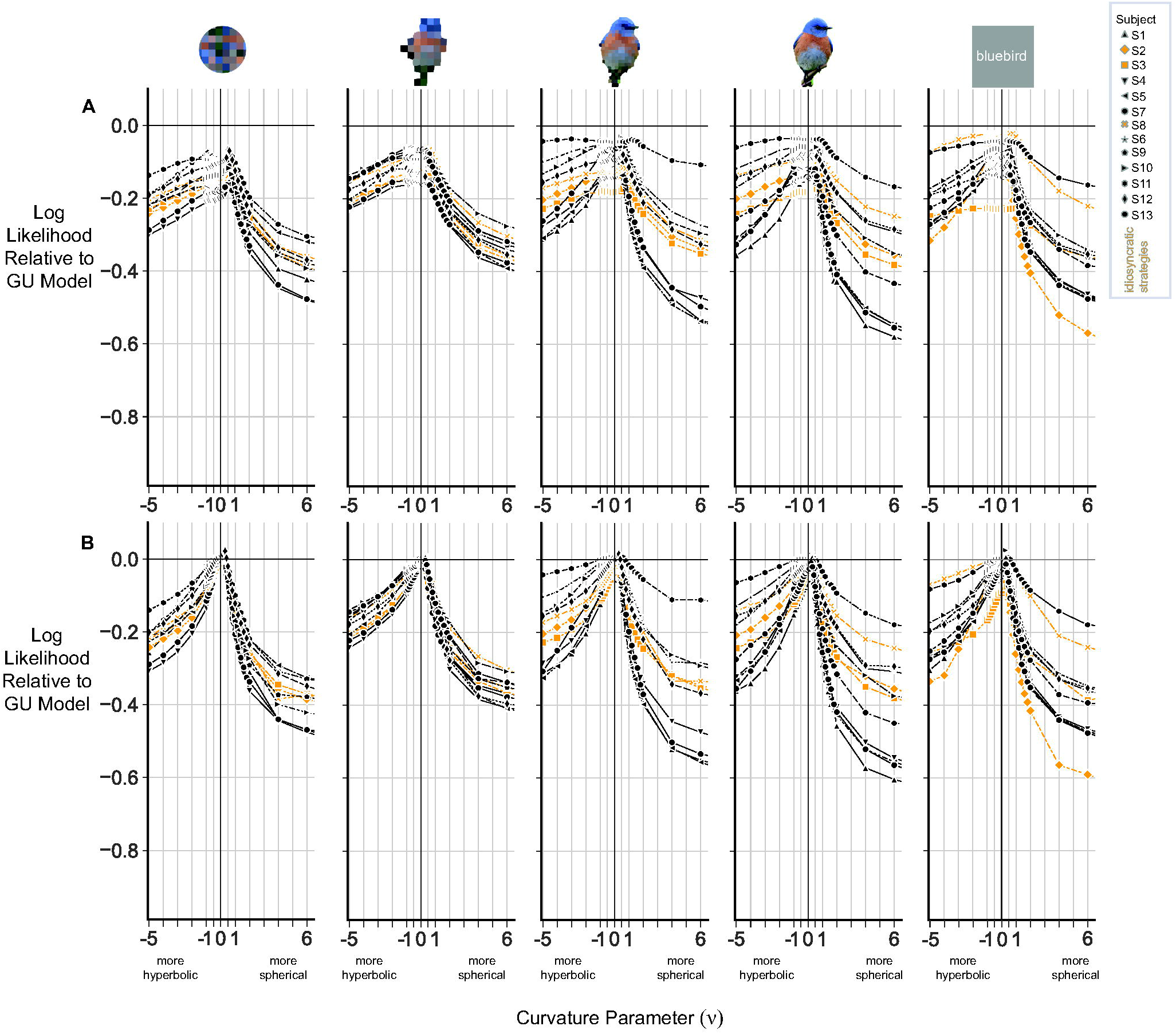
A small positive (spherical) curvature improves model fit. Model fit is quantified by the relative log likelihood per trial as in Figure 4. Negative values on the abscissa correspond to fits in a hyperbolic space(*v* = − *λ*, in eq. 7); positive values correspond to fits in a spherical space (*v* =2*μ*, from eq. 10). A positive value of *v* corresponds to the surface of a sphere with a radius of 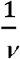 JNDs. The best model fit was typically found for *v* =0.2. **A:** 2-dimensional model. **B:** 7-dimensional model.

### Does the arrangement of points in a perceptual space vary across domains?

Having found no substantial differences in the geometric properties of the domains at the level of dimensionality or global curvature, we took a closer look at the spatial organization of points within each domain. As described below, we observed systematic shifts in the arrangements of points across domains. Characterizing these shifts identified differences in the representation of low-level domains and the domains with greater semantic content.

To visualize the arrangement of points in each domain, we projected the coordinates of points in the 5-dimensional model onto its first two principal components. The first two PCs explained between 45% and 81.16% (mean=60.6%) of the variance, across all participants and domains. Figure 6 shows the perceptual spaces for each participant (along rows) and domain (along columns).

**6.**
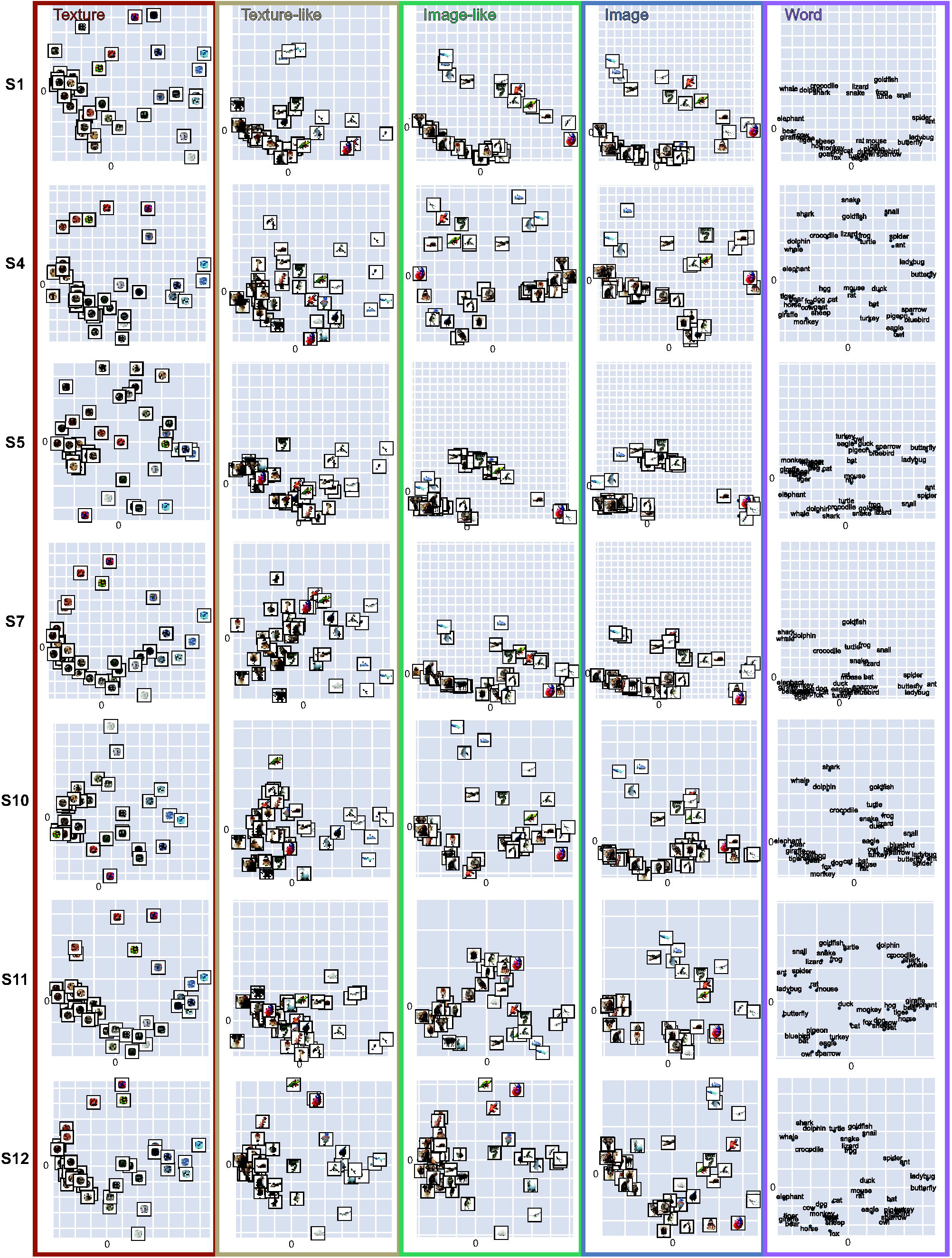
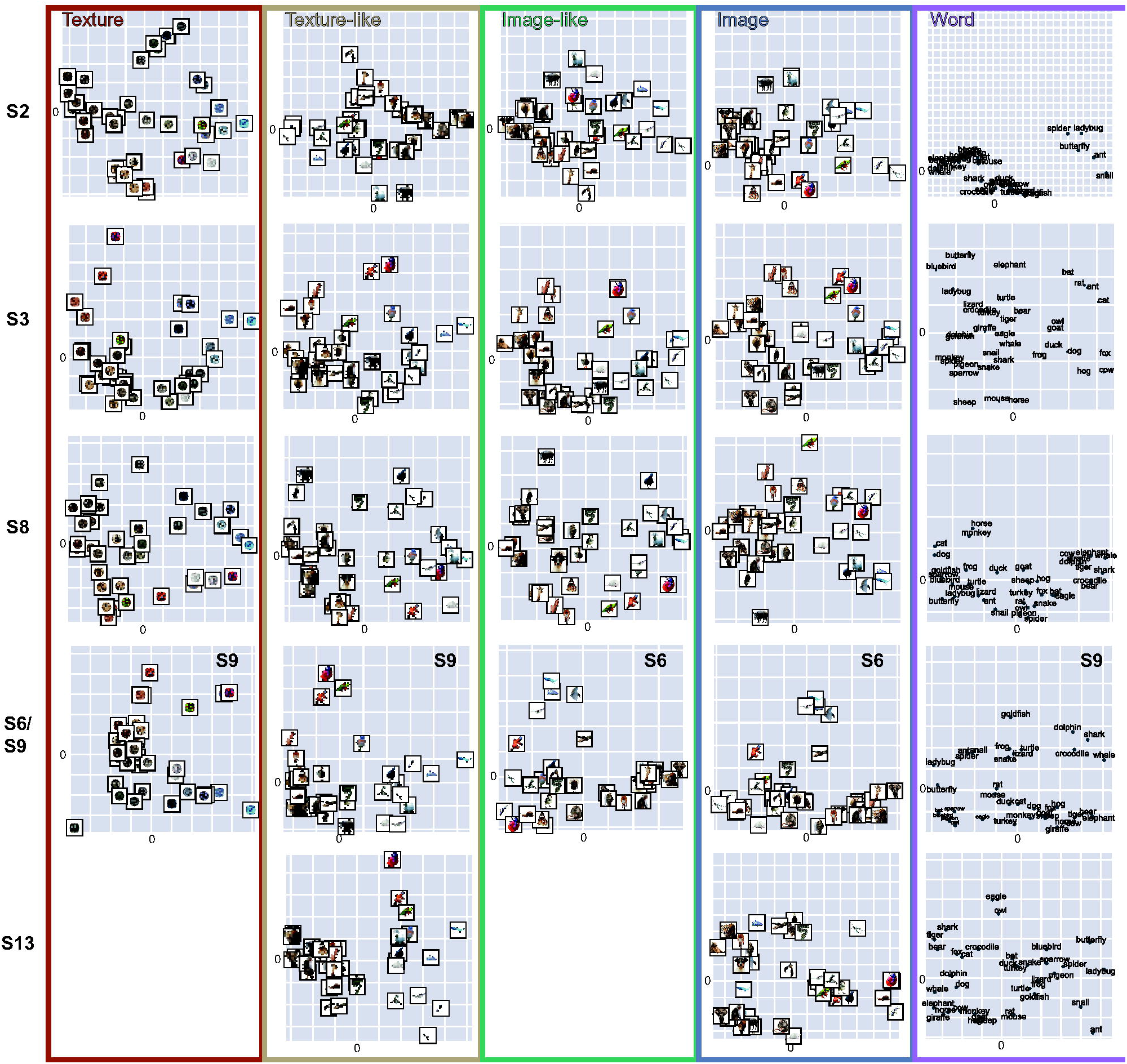
Arrangement of stimulus points in each of the five perceptual spaces (columns) for each participant. Scatterplots illustrate the projection of the 5-dimensional Euclidean coordinates onto the first two principal components. Lengths are in units of *σ*, the noticeable differences (JNDs), which is the additive noise in the decision model (eq. 1). Each scatterplot represents a given participant (row) and domain (column). **A:** Data from the participants S1, S4, S5, S7, S10, S11 and S12 who completed all five experiments. The data from participant S7 in the word domain were previously presented in (Waraich and Victor, 2022, Figure 6A). **B:** Data from participants who did not complete all experiments (S6, S9 and S13) and from participants with idiosyncratic strategies (S2, S3 and S8, see ‘Exclusion Criteria’).

A comparison of the arrangement of the 5-dimensional points across the domains within individual participants suggests a systematic change: in the word domain (see especially S1, S7, and S10 in the last column of Figure 6A), points appear to be grouped into arcs and clusters to a greater extent than in the texture and intermediate domains. The arrangements have intermediate characteristics as the domains transition from textures to words.

These differences reflect intrinsic relationships of the points and their distances and are not consequences of projecting the points into two dimensions. To demonstrate this, we calculated the Procrustes distance between the five domain configurations within each participant, a measure that quantifies how different two sets of points are from each other, after the best alignment that includes translation, rotation, and scaling (Figure 7). This analysis shows that for most participants, the Procrustes distance is high between the texture domain and all others, and between the texture-like domain and all others. However, the Procrustes distance between the image-like, image, and word domains is small for most participants, meaning that the points in these domains are arranged similarly. These shifts between the feature-weighted (texture- and texture-like) and semantic-weighted (image-like, image, and word) domains are apparent for most participants. For the outlier participants – who used idiosyncratic strategies for the word domain, the first four domains are all similar and the word domain stands out as different.

**7.**
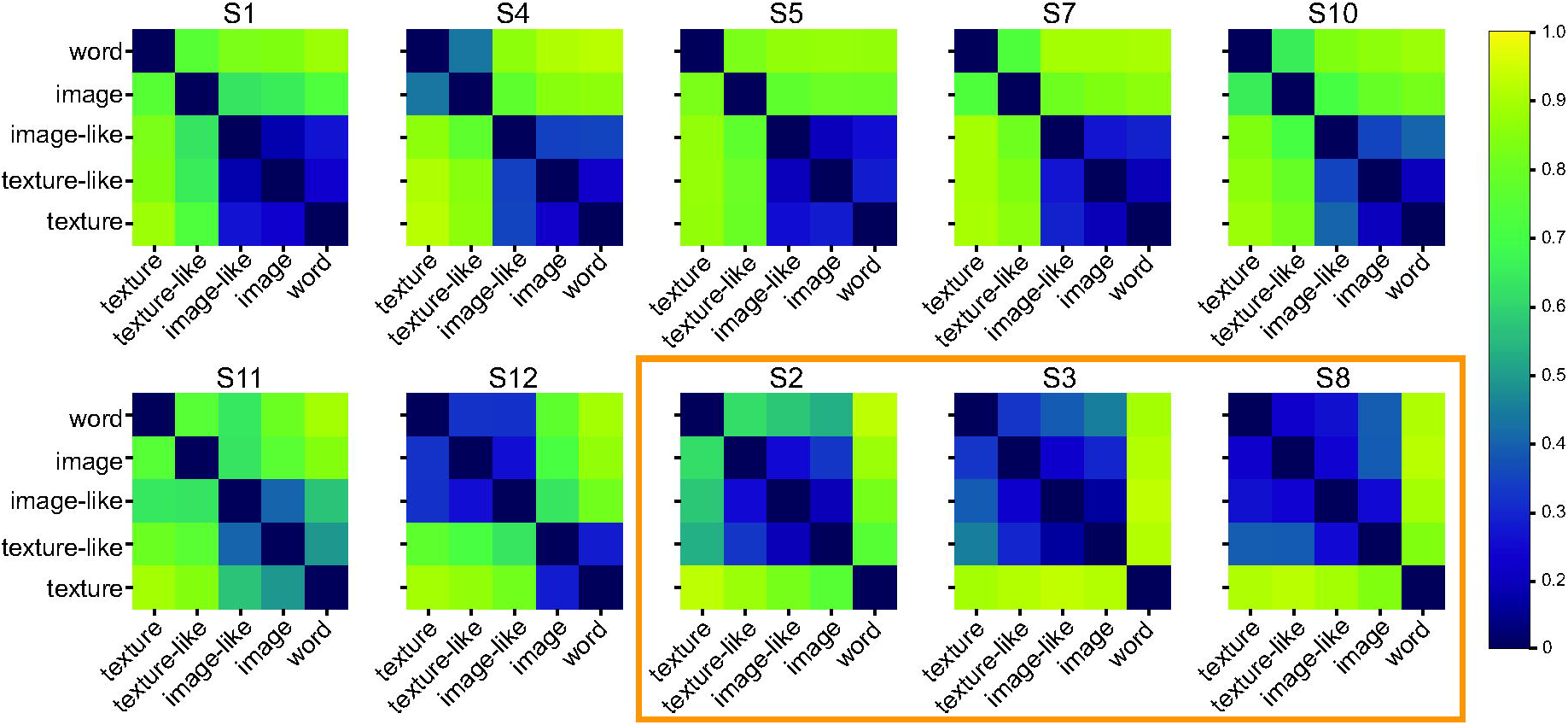
Comparison of the arrangement of points across domains, within each participant. Each heatmap shows the Procrustes distance (the fraction of the variance unexplained by the best coordinate rotation, reflection, and scaling) between each pair of domains; 0 indicates exact correspondence. The orange rectangle indicates the three participants with idiosyncratic strategies.

The coordinates deduced by the modeling procedure allow for an inference of the “size” of each space, in that the coordinates are all in units of JND’s, i.e., multiples of the internal noise parameter *σ*. Sizes (calculated as the root-mean-square distance between all points in a space, relative to *σ*) did vary somewhat (texture domain: 4.4-8.0 JND’s, texture-like domain: 4.0-5.9 JND’s, image-like domain: 2.3-9.7 JND’s, image domain: 2.8-10.6 JND’s, and word domain: 2.9-7.4 JND’s), but there were no consistent trends across participants or domains.

This range of values determines the sense in which the estimated curvature (Figure 5B) is small. The typical best-fit curvature *v* =0.2 corresponds to a sphere of radius 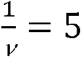 JND’s. Thus, the typical distances between points in the spherical space is much less than one half a great circle in that space 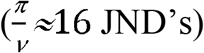.

### Do semantic-weighted domains have more tree-like perceptual spaces than low-level domains?

Finally, we directly tested the hypothesis that the arrangement of points in the word domain is more tree-like than in the feature domains – even if they have the same dimensionality. To do this, we used hierarchical clustering on coordinates of the 5-dimensional models, to generate dendrograms (see Figure 8A).

**8.**
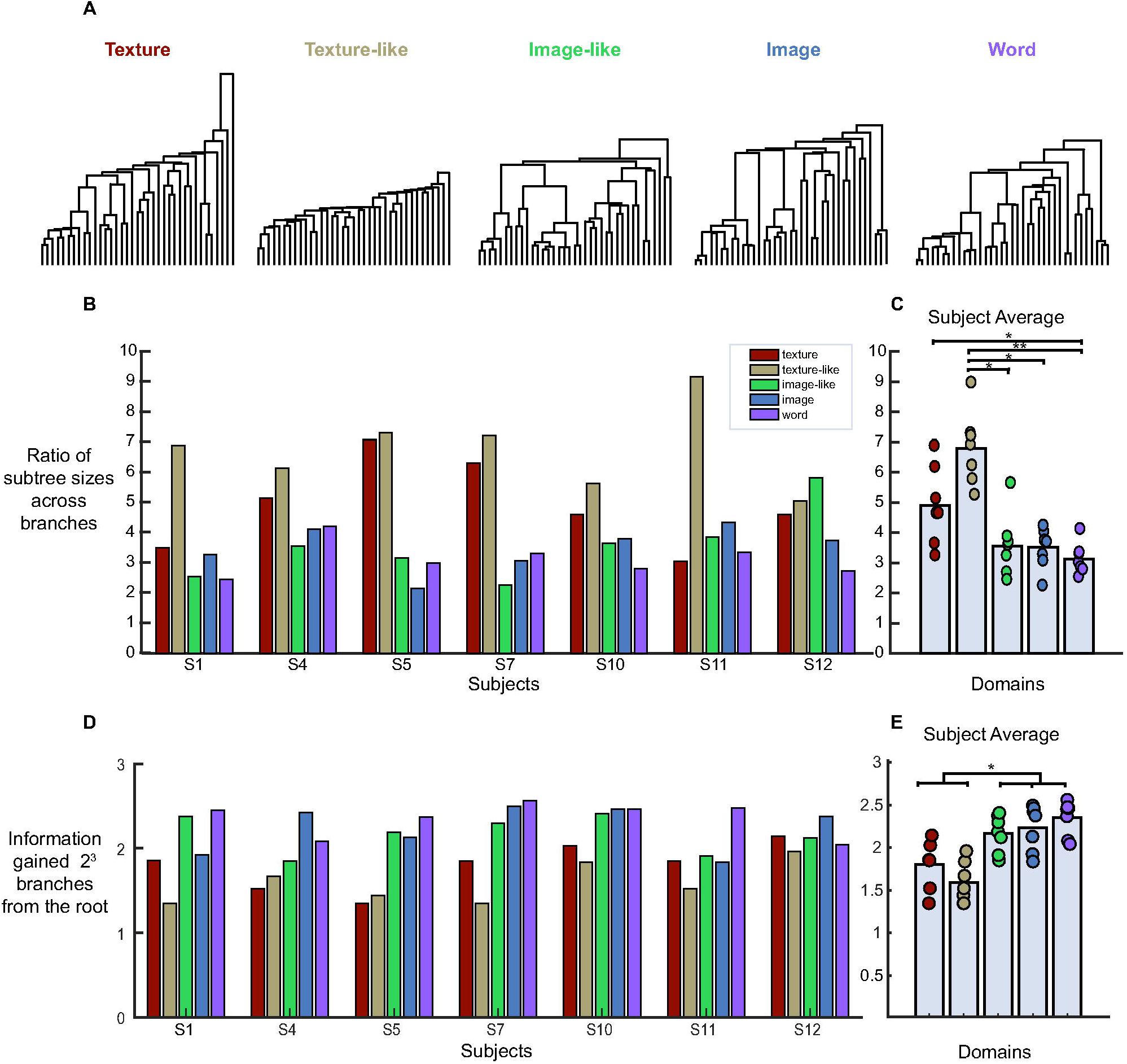
**A:** Dendrograms obtained from a representative participant (S7). Dendrograms were obtained from hierarchical clustering of the 5-dimensional coordinates for each domain’s perceptual space. **B:** The degree to which the branching structure of the dendrograms was balanced across levels of the tree, for all participants and domains. Balance is quantified by the mean ratio of the size of the larger subtree to the smaller subtree at each node; a ratio of 1 corresponds to maximal balance and larger values indicate more imbalance. **C:** Participant-wise average of **B**. Statistically significant differences indicated by *: p< 0.05 and **: p< 0.01 (paired t-tests, two-tailed, FDR-corrected). **D**: Information gained (in bits) after 2^3^ branches from the root of the tree, across participants and domains. **E:** Participant-wise average of **D**. Statistics as in **C**, significance at p<0.05 for differences between the texture domain and all three semantic domains (image-like, image and word) and between the texture-like domain and the three semantic domains (p <0.05 for all, p<0.01 for the difference between the texture-like domain and image domain).

We then used two measures to quantify how closely these dendrograms resembled a balanced binary tree. The first measure was the ratio of the subtree sizes across all branches of the tree; this has a minimum value of 1 for a balanced tree, and higher values indicate more unbalanced branches. The second measure captures how rapidly information is gained as one progresses from the root of the tree to its terminal branches: this is larger for a balanced tree than for an unbalanced one. Specifically, we compared the information gained 3 levels down, or after 2^3^ branches (this is approximately where the difference in information between a completely balanced binary tree and a completely unbalanced tree of 37 nodes would be maximal). Figures 8B-C and 8D-E respectively show these two measures for all participants and domains. The domains with more semantic information (image-like domain, image domain and word domain), yield trees that are more balanced in their branching structure than the texture-like tree, and there is a difference between the texture domain trees and word trees as well. Additionally, after traversing halfway down the trees, the trees derived from the more semantic domains (image-like, image and word) carry between 0.5 and 1 bit more of information, relative to trees derived from the low-level features (texture and texture-like domain. Overall, these results show that the geometry of the perceptual space shifts to a more tree-like representation as semantic information increases.

## Discussion

In this study, we asked whether low-level features and semantic information are represented in different ways. We hypothesized that, given the differences between the two kinds of information, the geometric characteristics of their mental representations would be distinct, specifically that the perceptual space for semantic information would have hierarchical characteristics. To address this question, we examined stimuli drawn from domains that carried a range of blends of semantic and feature-based information. Textures and texture-like stimuli varied in low-level visual features, e.g., color. Image-like stimuli, composed of readily recognizable pixelated pictures, contained more fine-grained shape and color information, and the image domain provided clear, immediately recognizable pictures, with all fine-grained structure of shape, texture and color preserved. Finally, the word domain – animal names written in lower-case – prompted the participants to think abstractly rather than attend to visual features. Participants viewed the domains in a pseudorandom order in which the higher-level domains were shown after the partially-texturized domains.

We found that the perceptual spaces of the lower- and higher-level domains were indeed different: the spaces with greater semantic content, (i.e., image-like, image and word domains) were more tree-like. The shift was rather sudden: the two low-level feature domains were similar to each other and less tree-like, whereas the image-like, image and word domains were all significantly more tree-like (see Figures 7, 8C and E), and with no significant differences between the image and word domains. Notably, this difference was seen across experiments that were matched for the number of stimuli, the paradigm, the instructions provided to participants, and the analysis method.

However, the differences in geometry across these domains was more subtle than one might have expected: there were no fundamental differences in the geometric characteristics of the perceptual spaces, e.g., their dimensionality or curvature.

Our study is complementary to previous ones (Huth et al., 2012; Hebart et al., 2020, 2023) that have used thousands of stimuli to probe the dimensionality of the perceptual space of objects. Huth et al. (2012) found at least 6-8 salient semantic dimensions by comparing fMRI activity patterns; Hebart et al. (2020) found 49, and later 66 (Hebart et al., 2023) independent, interpretable dimensions by analyzing similarity judgments. Hebart et al. (2020), in agreement with the literature (Caramazza and Shelton, 1998), find that one of these dimensions is animacy, and animals project onto this axis but also occupy other dimensions. Our stimuli likely probe this sector of the space and place a lower bound on the number of dimensions needed to represent it.

The new methodology used here allows us to add substantially to this picture, by specifying the geometry of this space and its relationship to the geometry to other perceptual spaces. Since we focus on a smaller number of objects (37 animals, vs. over 1000 objects), we probe the space in much greater detail: by collecting similarity judgments for ∼26% of possible triads, in contrast to the 0.05-0.15% coverage of triads in (Hebart et al., 2020, 2023). Because the current design includes repeated presentation of the triads, both in repeated and non-repeated contexts, we can estimate choice probability and context effects. The estimates of choice probability enable a maximum-likelihood modeling procedure based on a simple decision model, and as a result, the inferred perceptual spaces have interpretable units – JND’s . Global curvature is then determined in corresponding units (1/JND’s). Finally, there is sufficient data within individual participants to determine participant-specific geometry and consistency across participants.

Motivated in part by recent findings in olfaction (Zhou et al., 2018), we tested the hypothesis that hyperbolic spaces might provide a better model for perceptual spaces than Euclidean ones, as they can accommodate tree-like structures with less distortion than Euclidean models with the same number of dimensions (De Sa et al., 2018; Sonthalia and Gilbert, 2020). Instead, we found that all the domains could be embedded in a flat (Euclidean) or in a space with a modest amount of spherical (positive) curvature, as well as or better than hyperbolic (negatively curved) alternatives. Adding dimensions led to a greater improvement in modeling the perceptual space geometry than changing intrinsic curvature. However, in addition to the obvious difference between the domains (vision and olfaction), many other differences may contribute to the disparate findings. A key procedural difference is that we compared similarities directly, while Zhou et al. (2018) inferred them from a model based on feature vectors. A key analytic difference is that we constructed geometric models via a maximum-likelihood approach; Zhou et al. (2018) inferred global geometry via a topological approach that determined persistent homologies.

Given that the higher-level spaces are more tree-like, and also that hyperbolic spaces are especially suitable for embedding trees (De Sa et al., 2018; Keller-Ressel and Nargang, 2020; Matsumoto et al., 2021), it may appear surprising that we don’t find indications of negative curvature in the higher-level spaces. However, this finding likely reflects two factors. First, although the volume of a hyperbolic sphere indeed grows exponentially with radius–as is appropriate for embedding trees, since the number of tree nodes grows exponentially with the number of generations – embedding a tree in a low-dimensional hyperbolic space nevertheless leads to a distortion of the tree distances. For example, a node should have identical tree distances to any other node with a different parent but the same grandparent (i.e., to all of its first-cousins), but the hyperbolic distances from one node to its leftmost and rightmost first-cousins differ greatly. To cure this problem, one needs an embedding with one dimension for each level of the tree (see, for example, (Saxe et al., 2019) Fig. 9 lower row), and this embedding can be Euclidean. The second consideration is that the exponential growth in volume that characterizes hyperbolic geometry is accompanied by an exponential growth in hyperbolic distance between points at a given radius. This behavior is likely inconsistent with our data, and would make a hyperbolic geometry a poor fit.

Our characterization of perceptual space geometry is a global one: it assumes uniformity across the space and across scales, and this is almost certainly a simplification. A hybrid of a tree-representation at large scales and a continuous one at small scales is both consistent with our data and makes intuitive sense. For example, one key feature that many participants used was size, a continuous variable. This suggests that the semantic domains may be tree-like for coarse distinctions (e.g., air vs. land vs. sea animal), and continuous at finer scales.

Although a more precise characterization of geometry may add nuance to the global characterization used here, our main finding is that the shift in the tree-likeness of the domains is relatively abrupt. Within the semantic domains, we did not find any statistically significant difference in the geometry of their representations. The perceptual learning literature (Hochstein and Ahissar, 2002) suggests a possible explanation: that in the semantic domains, the ‘first conscious percept’ that arises in the mind is the gist of the scene, i.e., the animal, and once an animal is recognized, it can’t be unseen. If the gist is more important than variations in lower-level features, there may not be any stagewise, slow changes in perceptual geometry at all, but rather a jump at the point where the stimulus becomes meaningful to the observer.

## Acknowledgments

We thank our participants for the time they devoted to the experiments. We thank Martin Hebart, Laurence T. Maloney, Nikolaus Kriegeskorte, Shaul Hochstein, and Mary M. Conte for their helpful discussions. A portion of this work was presented as posters at meetings of the Vision Sciences Society (2021, 2022) and the Society for Neuroscience (2022). The work was funded by EY07977 to JV and the Fred Plum Fellowship in Systems Neurology and Neuroscience.

